# Effects of nonantibiotic alternative growth promoter combinations on nutrient utilization, digestive enzyme activity, intestinal morphology, and cecal microflora of broilers

**DOI:** 10.1101/2022.12.20.521250

**Authors:** Zunyan Li, Beibei Zhang, Weimin Zhu, Yingting Lin, Jia Chen, Fenghua Zhu, Yixuan Guo

## Abstract

Given the ban on antibiotic growth promoters, six nonantibiotic alternative growth promoter combinations (NAGPCs) for broilers were evaluated. All birds were fed pellets of two basal diets—starter (0−21 d) and grower (22−42 d)—with either enramycin (ENR) or NAGPC supplemented. 1) control + 100 mg/kg ENR; 2) control diet (CON, basal diet); 3) control + 2,000 mg/kg mannose oligosaccharide (MOS) + 300 mg/kg mannanase (MAN) + 1,500 mg/kg sodium butyrate (SB) (MMS); 4) control + 2,000 mg/kg MOS + 300 mg/kg MAN + 500 mg/kg *Bacillus subtilis* (BS) (MMB); 5) control + 2,000 mg/kg MOS + 9,000 mg/kg fruit oligosaccharide (FOS) + 1,500 mg/kg SB (MFS); 6) control + 9,000 mg/kg FOS + 500 mg/kg BS (MFS) (MBP). The experiment used a completely random block design with six replicates per group: 50 Ross 308 broilers in the starter phase and 16 in the grower phase. All the NAGPCs significantly improved (P < 0.01) utilization of dry matter (DM), organic matter (OM), crude protein (CP), and crude fat (CF) on d 21, significantly increased DM, OM (P < 0.01), and CP (P < 0.05) on d 42, and significantly increased (P < 0.01) villus height, crypt depth, and the villus height to crypt depth ratio in the jejunum and ileum compared with CON and ENR. On d 21 and 42, trypsin, lipase, and amylase activity of the duodenum significantly increased in the MMS, MMB, MFB, and MFM groups. Compared with ENR and CON, MMS, MMB, and MBP increased the abundance of *Firmicutes* at d 21 and of *Bacteroides* at d 42 whereas MMB, MFB, and MBP decreased the abundance of *Proteobacteria* at d 21 and 42. Overall, the NAGPCs were found to have some beneficial effects and may be used as effective antibiotic replacements in broilers.

## Introduction

Antibiotics growth promoters (AGPs) are used in broiler diets to manage illnesses, maintain health, stimulate growth, and enhance feed utilization. However, the threats that medication resistance and antibiotic residues pose to broiler and human health have raised significant alarm regarding AGP use [1]. After the European Union and United States prohibited the use of AGPs, China followed suit in 2020. The prohibition on AGP use prompted the industry to develop suitable antibiotic replacements [2]. Growth rate reduction induced by the withdrawal of AGPs can reduce production efficiency as well as impact food safety and broiler health [3]. Therefore, in the absence of AGPs, alternate methods need to be developed to ensure feed efficiency and broiler health [4].

Currently, nonantibiotic alternative growth promoters (NAGPs) such as enzymes, probiotics, prebiotics, and acidifiers are being used in broiler feed to replace antibiotics [5]. As the energy source of intestinal bacteria, butyric acid in sodium butyrate (SB) can improve the intestinal flora structure and nutrient digestibility [6]. The digestive capacity of the intestinal tract can be improved using the digestive enzymes released by *Bacillus subtilis* (BS) [7]. By preventing colonization of intestinal pathogenic bacteria, prebiotics dominated by mannose oligosaccharide (MOS) can also improve the intestinal environment [8]. Mannanase (MAN) can hydrolyze the non-starch polysaccharide present in soya bean meal and supplement endogenous enzymes to increase nutrient digestibility [9]. Owing to its unique spatial structure, phytase (PT) can breakdown phytic acid into inositol and inorganic phosphorus and promote the release of other nutrients along with phytic acid [10]. The different mechanisms and modes of action of these enzymes, probiotics, prebiotics, and organic acids suggest the potential for complementary synergistic effects upon supplementing diets with a mixture of these additives.

In a previous study comparing the effects of formic acid, propionic acid, and their combination on the growth performance of broilers, the feed conversion ratio for only formic acid + propionic acid dropped significantly compared with that for the other combinations [11]. Another study on poultry diets showed that a combination of xylanase, amylase, and protease could elicit greater improvements in nutrient utilization than xylanase, amylase, or protease given alone [12]. These results indicate that NAGPCs have better effect than NAGPs alone. However, current research on NAGPSs is mainly focused on the effect of combining the same kinds of NAGPs in broilers. The beneficial effect of synergy may be limited by the combination of the same kind of NAGPs.

Therefore, the objective of this study was to evaluate the effects of six NAGPCs with different types of NAGPs on nutrient utilization, digestive enzyme activity, intestinal morphology, and cecal microflora of broilers so as to find a better alternative method to ensure feed efficiency and the broiler health.

## Materials and methods

### Ethics approval

Animal care and experimental protocols (License no. QAU20210607) were approved by the Animal Care and Use Committee of Qingdao Agricultural University, China. The animals were cared for according to the Animal Care Guidelines of China. All efforts were made to minimize animal suffering.

### Birds and housing

In the starter phases, 2,400 1-d-old healthy Ross 308 broilers of both sexes with similar body weight (BW) were randomly assigned to eight groups; each group included six repeats with 50 chickens each. In the grower phases, 768 21-d-old healthy Ross 308 broilers with similar BW were randomly assigned to eight groups; each group included six repeats with 16 chickens each. The feeding experiment was conducted for 21 d after individual weighing. No significant differences in the initial BW were observed among these groups. Seven-d-old chickens were immunized with live Newcastle disease and infectious bronchitis vaccines by nose and eye dripping. Fourteen-d-old chickens were immunized with a live infectious bursal disease vaccine in drinking water. At 21 d of age, chickens were immunized with a live Newcastle disease vaccine by nose and eye dripping. During the early stages of the experiment, 50 broilers in each replicate were reared in a single cage (130 cm × 55 cm × 50 cm). During the latter stages of the experiment, each replicate was housed in cages with 16 birds per cage. The broilers had free access to pellet feed and water throughout the study. In a controlled environment with a relative humidity of 45–55% and temperature of 25– 34 °C, the broilers were maintained under an 18 h/6 h light/dark cycle. In the first week of the experiment, the ambient temperature was maintained at 34 °C; it was then gradually decreased to 26 °C after 21 d and maintained thereafter. The ingredients and nutrient levels of the diets are listed in Table 1.

**Table 1.** Ingredient (%) and nutritive value of a basal diet.

| Item | Starter feed | Grower feed |
| --- | --- | --- |
| Ingredient (%) |  |  |
| Corn | 45.00 | 44.30 |
| Wheat | 15.00 | 14.70 |
| Expanded soybean meal | 23.50 | 22.00 |
| Cottonseed meal | 5.00 | 5.00 |
| Corn gluten meal | 2.50 | 1.00 |
| Hydrolyzed feather meal | 1.50 | 1.00 |
| Calcium bicarbonate | 0.80 | 0.60 |
| Stone powder | 1.70 | 1.70 |
| Bentonite | 0.50 | 0.50 |
| Soybean oil | 2.20 | 7.00 |
| Sodium chloride | 0.25 | 0.25 |
| Lysine | 0.65 | 0.62 |
| Methionine | 0.25 | 0.23 |
| Threonine | 0.15 | 0.10 |
| Premix<sup>1</sup> | 1.00 | 1.00 |
| Total | 100.00 | 100.00 |
| Nutrition level<sup>2</sup> |  |  |
| ME (kcal/kg) | 2900 | 3200 |
| CP (%) | 22.00 | 20.00 |
| Calcium (%) | 0.98 | 0.88 |
| Total phosphorus (%) | 0.76 | 0.65 |
| Lysine (%) | 1.41 | 1.37 |
| Methionine (%) | 0.56 | 0.51 |
| Threonine (%) | 0.91 | 0.80 |

<sup>1</sup>Vitamin-mineral premix (each kg contained); vitamin A, 9050 IU; vitamin D3, 1950 IU; vitamin E, 26 IU; vitamin K3, 5.0 mg; vitamin B1, 2.6 mg; vitamin B2, 8.0 mg; pantothenic acid, 15.0 mg; vitamin B6,3.0 mg; vitamin B12, 0.02 mg; nicotinic acid, 35 mg; choline chloride, 9050 mg; biotin, 0.20 mg; folic acid, 1.2 mg; Mn,60 mg. Fe,80 mg; Zn,60 mg; Cu,8.5 mg; I,0.27 mg; Se,0.20 mg <sup>2</sup>Nutrient levels were calculated by analysis.

### Experimental design

The amounts of different NAGPs and NAGPCs to be added were selected based on the previous *in vitro* digestion test. All birds were fed pellets of two basal diets (Table 1)—starter (0−21 d) or grower (22−42 d)—that differed within each phase by the addition of either enramycin (ENR) or NAGPCs, as follows: 1) control + 100 mg/kg ENR; 2) a control diet (CON, basal diet); 3) control + 2,000 mg/kg MOS + 300 mg/kg MAN + 1,500 mg/kg SB (MMS); 4) control + 2,000 mg/kg MOS + 300 mg/kg MAN + 500 mg/kg BS (MMB); 5) control + 2,000 mg/kg MOS + 9,000 mg/kg FOS + 1,500 mg/kg SB (MFS); 6) control + 2,000 mg/kg MOS + 9,000 mg/kg FOS + 500 mg/kg BS (MFB); 7) control + 2,000 mg/kg MOS + 9,000 mg/kg FOS + 300 mg/kg MAN (MFM); and 8) control + 2,000 mg/kg MOS + 500 mg/kg BS + 37 mg/kg PT (MBP). SB (≥ 50%) was obtained from Qilu Animal Health Co., Ltd. MOS (≥ 12%) was obtained from Orteki Biological products Co., Ltd. Fruit oligosaccharide (FOS, ≥ 95%) was obtained from Baolingbao Biological Co., Ltd. MAN (≥ 50,000 U/g) was obtained from Shandong Shengdao Biotechnology Co., Ltd. PT (≥ 3 × 10^5^ U/g) was obtained from Jinan Baisijie Biological Engineering Co., Ltd. BS (≥ 2 × 10^11^ cfu/g) was obtained from Beijing Keweibo Biotechnology Co., Ltd. The products were added manually, and no specific premix or carrier was used. The experimental materials were all products available on the market.

### Sample collection

#### Nutrient utilization

Each dietary phase began with providing a 500 g sample of feed. Excreta were collected daily from d 19 to 21 and from d 40 to 42, pooled in a cage on the last day of collection, mixed well, and subsampled. The subsamples were then lyophilized, ground, and passed through a 0.5 mm screen. Excreta and diet samples were analyzed for dry matter (DM) (AOAC Official method 930.15, 2006) [13], ash (AOAC Official method 942.05, 2006) [13], crude protein (CP) (AOAC Official method 990.02, 1990) [14], ether extract (AOAC Official method 968.06, 2000) [15], calcium (Ca) and phosphorus (P) (AOAC Official method 2011.14, 1990) [14], and acid insoluble ash (AIA) (AOAC Official method 975.12, 1990) [14]. The ether extract was crude fat (CF). Nutrient utilization was determined using AIA as an indicator. Feed and excreta were corrected based on DM. Nutrient utilization was calculated from the following equation:

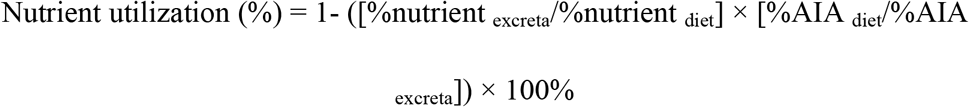

where nutrient indicates the nutrient content in feed or excreta, and AIA indicates acid insoluble ash content in feed or excreta.

#### Measurement of digestive enzyme activity

At d 21 and 42, one broiler from each replicate (six per treatment) was euthanized by cervical dislocation. One gram of duodenal content was removed using an aseptic scraper and placed in a 2 mL aseptic tube that was frozen at −20°C. Amylase, lipase, and trypsin activity in the duodenum was determined using corresponding diagnostic kits (Nanjing Jiancheng Bioengineering Institute, Nanjing, China) according to the manufacturer’s instructions.

#### Villus histomorphometry

Following euthanasia, the mid-jejunum and mid-ileum segments were removed (about 2 cm). After emptying their contents with distilled water using a Nalgene LDPE wash bottle, each tissue was individually fixed in a formalin solution and stored at room temperature. The jejunum was then used to measure the intestinal villi and crypts. The small intestine samples (jejunum and ileum) were fixed in 4% buffered formaldehyde and then dehydrated using a graded series of xylene and ethanol before embedding in paraffin for histological analysis. The small intestine sections (8 microns in length) were then deparaffinized using xylene and rehydrated using graded ethanol dilutions. Hematoxylin and eosin were used to stain the slides. Ten slides were prepared for each sample (from the central region of the sample), and images were captured using an optical binocular microscope (OLYMPUS CKX53, Jingkai instrument and equipment Co., Ltd., Suzhou, China). The villus height and crypt depth were measured by Image – Pro Plus 6. 0 five times from distinct villi and crypts on each slide. Averages were computed for these values.

#### 16S rDNA sequencing and data analysis

After euthanasia, one gram of cecal content was removed using an aseptic scraper and placed in a 2 mL aseptic tube that was frozen at −20°C. Total genomic DNA from cecal digesta was extracted using PowerSoil® DNA Isolation Kit (Sigma-Aldrich, Carlsbad, United States of America). DNA concentration and purity were assessed using a Synergy HTX (Gene Company Limited, Shanghai, China). DNA amplicons from individual samples were amplified using polymerase chain reaction with specific primers for the V3-V4 regions of the 16S rRNA gene. Amplicons generated from each sample were subjected to agarose (1%) gel electrophoresis, excised, purified using a Monarch DNA kit (Jizhi biochemical Technology Co., Ltd., Shanghai, China), and quantified using the Synergy HTX system. The constructed library was sequenced on the Illumina Novaseq6000 PE250 platform (Jingneng Biotechnology Co., Ltd., Shanghai, China). The original data were spliced (FLASH, version 1.2.11); the spliced sequences were filtered by quality (Trimmomatic, version 0.33), and chimerism (UCHIME, version 8.1) was removed to obtain high-quality tag sequences. The sequences were clustered at the level of 97% similarity using USEARCH (version 10.0). By default, 0.005% of all sequenced sequences was taken as the threshold for filtering operational taxonomic units (OTUs). Alpha diversity analysis was performed using a classifier Bayesian algorithm (https://qiime2.org/) and the QIIME 2 software. Principal coordinate analysis (PCoA) and beta diversity analysis implemented in the QIIME 2 software were performed based on UniFrac distance matrices. Linear discriminant analysis Effect Size (LEFSe) used Kruskal-Wallis rank sum to detect species with significant abundance differences between groups. The Wilcoxon rank-sum test was then used to determine the consistency of the differences between different subgroups. Finally, linear regression analysis (LDA) was used to estimate the magnitude of the influence of the abundance of each component (species) on the differential effect. Color correlograms were generated using the R corrplot package (Version 0.84).

### Statistical analyses

All data were analyzed using one-way analysis of variance (ANOVA) and the *post hoc* Duncan’s multiple range test using the SPSS statistical software (version 20.0; SPSS Inc., Chicago, IL, USA). P < 0.05 was considered to indicate statistical significance. The results are expressed as the mean and pooled standard error of mean (SEM).

## Results

### Nutrient utilization

On d 21, DM, OM, CP, and CF utilization increased significantly (P < 0.01) in the MMS, MMB, MFS, MFB, MFM, and MBP groups compared with that in the CON and ENR groups, whereas the treatments had no effect on Ca (P = 0.91) and P (P = 0.6) utilization (Table 2). On d 42, a significant increase was observed in DM, OM, and CP (P < 0.01) utilization in the MMS, MMB, MFS, MFB, MFM, and MBP groups compared with that in the CON and ENR groups as well as in CF utilization (P < 0.01) in the MMS, MMB, MFB, and MFM groups. On d 42, a significant increase in Ca (P = 0.04) and P (P < 0.01) utilization was found in the MMS, MMB, MFS, MFB, MFM, and MBP groups compared with that in the CON group.

**Table 2.**
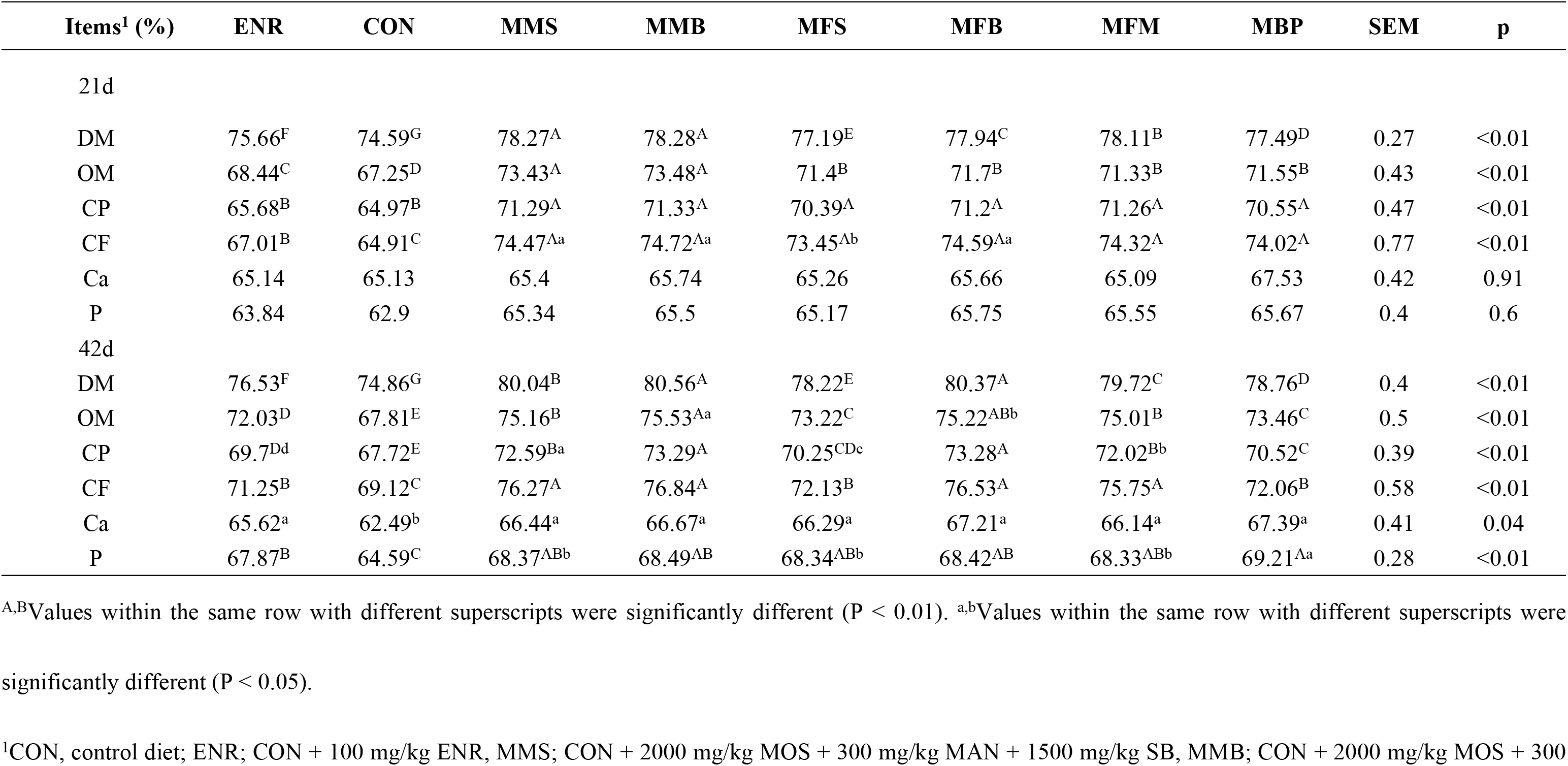
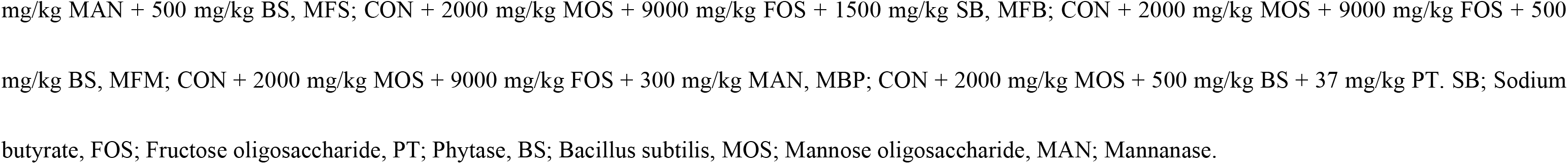
Effects of dietary nonantibiotic alternative growth promoter combinations on the utilization of dry matter (DM), organic matter (OM), crude protein (CP), crude fat (CF), calcium (Ca), and phosphorus (P) in broiler chicks at 21 and 42 days.

| Items<sup>1</sup> (%) | ENR | CON | MMS | MMB | MFS | MFB | MFM | MBP | SEM | p |
| --- | --- | --- | --- | --- | --- | --- | --- | --- | --- | --- |
| 21d |  |  |  |  |  |  |  |  |  |  |
| DM | 75.66<sup>F</sup> | 74.59<sup>G</sup> | 78.27<sup>A</sup> | 78.28<sup>A</sup> | 77.19<sup>E</sup> | 77.94<sup>C</sup> | 78.11<sup>B</sup> | 77.49<sup>D</sup> | 0.27 | <0.01 |
| OM | 68.44<sup>C</sup> | 67.25<sup>D</sup> | 73.43<sup>A</sup> | 73.48<sup>A</sup> | 71.4<sup>B</sup> | 71.7<sup>B</sup> | 71.33<sup>B</sup> | 71.55<sup>B</sup> | 0.43 | <0.01 |
| CP | 65.68<sup>B</sup> | 64.97<sup>B</sup> | 71.29<sup>A</sup> | 71.33<sup>A</sup> | 70.39<sup>A</sup> | 71.2<sup>A</sup> | 71.26<sup>A</sup> | 70.55<sup>A</sup> | 0.47 | <0.01 |
| CF | 67.01<sup>B</sup> | 64.91<sup>C</sup> | 74.47<sup>Aa</sup> | 74.72<sup>Aa</sup> | 73.45<sup>Ab</sup> | 74.59<sup>Aa</sup> | 74.32<sup>A</sup> | 74.02<sup>A</sup> | 0.77 | <0.01 |
| Ca | 65.14 | 65.13 | 65.4 | 65.74 | 65.26 | 65.66 | 65.09 | 67.53 | 0.42 | 0.91 |
| P | 63.84 | 62.9 | 65.34 | 65.5 | 65.17 | 65.75 | 65.55 | 65.67 | 0.4 | 0.6 |
| 42d |  |  |  |  |  |  |  |  |  |  |
| DM | 76.53<sup>F</sup> | 74.86<sup>G</sup> | 80.04<sup>B</sup> | 80.56<sup>A</sup> | 78.22<sup>E</sup> | 80.37<sup>A</sup> | 79.72<sup>C</sup> | 78.76<sup>D</sup> | 0.4 | <0.01 |
| OM | 72.03<sup>D</sup> | 67.81<sup>E</sup> | 75.16<sup>B</sup> | 75.53<sup>Aa</sup> | 73.22<sup>C</sup> | 75.22<sup>ABb</sup> | 75.01<sup>B</sup> | 73.46<sup>C</sup> | 0.5 | <0.01 |
| CP | 69.7<sup>Dd</sup> | 67.72<sup>E</sup> | 72.59<sup>Ba</sup> | 73.29<sup>A</sup> | 70.25<sup>CDc</sup> | 73.28<sup>A</sup> | 72.02<sup>Bb</sup> | 70.52<sup>C</sup> | 0.39 | <0.01 |
| CF | 71.25<sup>B</sup> | 69.12<sup>C</sup> | 76.27<sup>A</sup> | 76.84<sup>A</sup> | 72.13<sup>B</sup> | 76.53<sup>A</sup> | 75.75<sup>A</sup> | 72.06<sup>B</sup> | 0.58 | <0.01 |
| Ca | 65.62<sup>a</sup> | 62.49<sup>b</sup> | 66.44<sup>a</sup> | 66.67<sup>a</sup> | 66.29<sup>a</sup> | 67.21<sup>a</sup> | 66.14<sup>a</sup> | 67.39<sup>a</sup> | 0.41 | 0.04 |
| P | 67.87<sup>B</sup> | 64.59<sup>C</sup> | 68.37<sup>ABb</sup> | 68.49<sup>AB</sup> | 68.34<sup>ABb</sup> | 68.42<sup>AB</sup> | 68.33<sup>ABb</sup> | 69.21<sup>Aa</sup> | 0.28 | <0.01 |

<sup>A,B</sup>Values within the same row with different superscripts were significantly different (P < 0.01). <sup>a,b</sup>Values within the same row with different superscripts were significantly different (P < 0.05).
<sup>1</sup>CON, control diet; ENR; CON + 100 mg/kg ENR, MMS; CON + 2000 mg/kg MOS + 300 mg/kg MAN + 1500 mg/kg SB, MMB; CON + 2000 mg/kg MOS + 300

### Duodenal digestive enzyme activity

On d 21, a significant increase in duodenal trypsin, amylase, and lipase (P < 0.01) activity was observed in the MMS, MMB, MFS, MFB, MFM, and MBP groups compared with that the CON and ENR groups (Table 3). On d 42, a significant increase in the duodenal trypsin and lipase (P < 0.01) activity in the MMS, MMB, MFS, MFB, MFM, and MBP groups was observed compared with that in the CON and ENR groups; furthermore, a significant increase in duodenal amylase (P < 0.01) activity was observed in the MMS, MMB, MFB, and MFM groups.

**Table 3.**
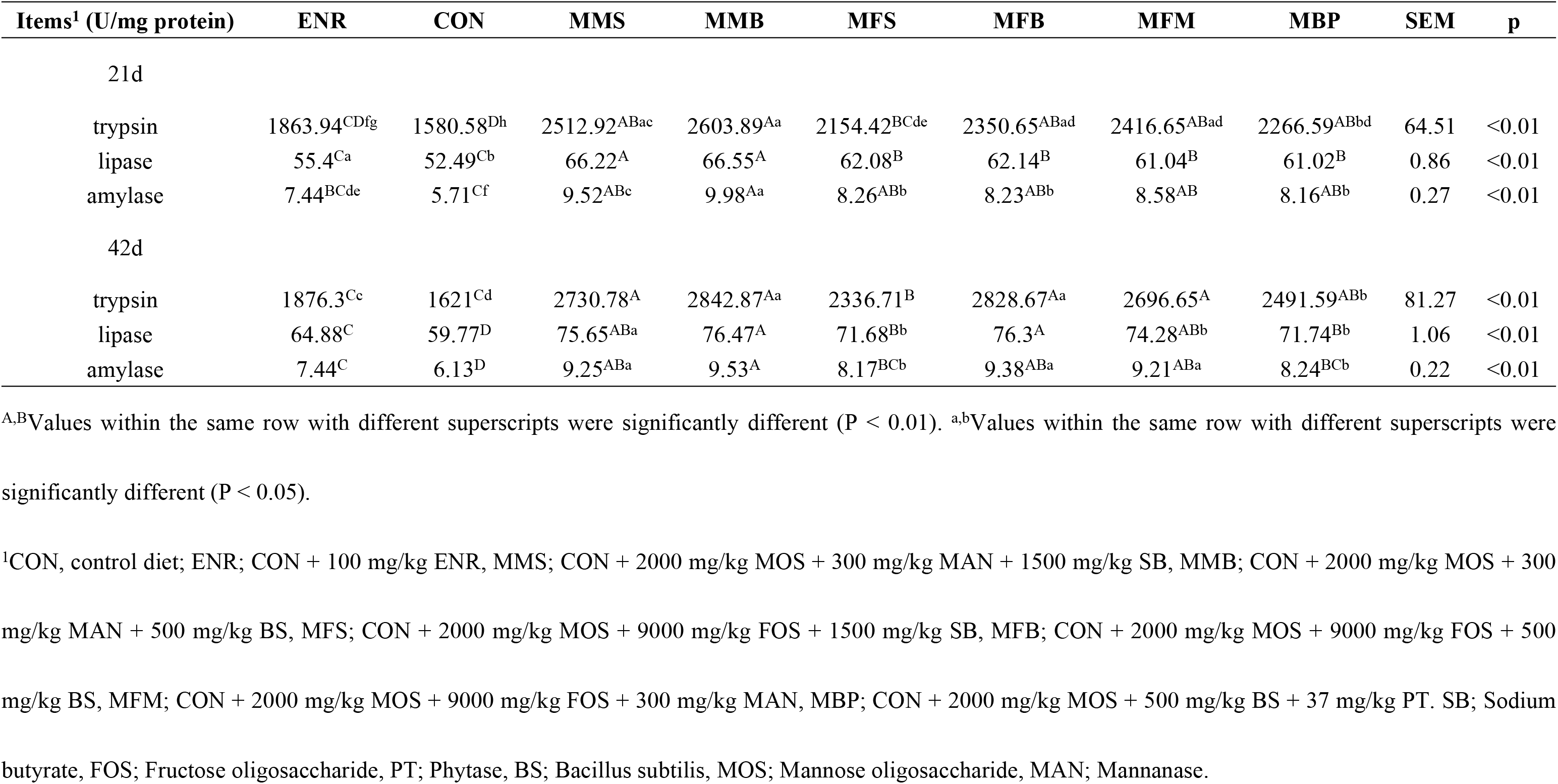
Effects of dietary nonantibiotic alternative growth promoter combinations on duodenal digestive enzyme activity of broilers at 21 and 42 days.

### Ileum and jejunum histology

On d 21, the villus height (P = 0.02) and villus height/crypt depth (P = 0.04) in the jejunum were significantly increased in the MMS, MMB, MFB, and MFM groups compared with those in the CON and ENR groups (Table 4, Figs 1, 2, 3, and 4). Compared with those in the CON and ENR groups, the ileal villus height (P = 0.03) and villus height/crypt depth (P < 0.01) were significantly increased and crypt depth (P = 0.01) was significant decreased in the MMS, MMB, MFS, MFB, MFM, and MBP groups. On d 42, the villus height (P = 0.01) and villus height/crypt depth (P < 0.01) in the jejunum were significant increased in the MMS, MMB, MFS, MFB, MFM, and MBP groups compared with those in the CON and ENR groups; crypt depth (P = 0.01) was also significantly decreased. Ileal villus height (P < 0.01) was significantly increased in the MMS, MMB, MFS, MFB, MFM, and MBP groups compared with that in the CON and ENR groups; furthermore, the crypt depth (P < 0.01) showed a significant decrease, whereas the villus height/crypt depth ratio (P < 0.01) showed a significant increase.

**Table 4.** Effects of dietary nonantibiotic alternative growth promoter combinations on ileal and jejunum histomorphological parameters of broilers at 21 and 42 days.

| Items<sup>1</sup> | ENR | CON | MMS | MMB | MFS | MFB | MFM | MBP | SEM | p |
| --- | --- | --- | --- | --- | --- | --- | --- | --- | --- | --- |
| 21d |  |  |  |  |  |  |  |  |  |  |
| jejunum |  |  |  |  |  |  |  |  |  |  |
| villus height (μm) | 562.21<sup>b</sup> | 320.43<sup>b</sup> | 922.07<sup>a</sup> | 939.13<sup>a</sup> | 681.10 | 911.74<sup>a</sup> | 905.55<sup>a</sup> | 690.28 | 55.33 | 0.02 |
| crypt depth (μm) | 105.18 | 118.22 | 93.30 | 92.95 | 96.05 | 93.54 | 94.18 | 96.54 | 3.32 | 0.57 |
| villus height/crypt depth | 5.47<sup>b</sup> | 3.07<sup>b</sup> | 10.07<sup>a</sup> | 10.19<sup>a</sup> | 7.33 | 9.84<sup>a</sup> | 9.61<sup>a</sup> | 7.52 | 0.67 | 0.04 |
| ileal |  |  |  |  |  |  |  |  |  |  |
| villus height (μm) | 486.82<sup>b</sup> | 435.45<sup>b</sup> | 845.27<sup>a</sup> | 873.05<sup>a</sup> | 792.17<sup>a</sup> | 827.22<sup>a</sup> | 812.34<sup>a</sup> | 805.29<sup>a</sup> | 44.15 | 0.03 |
| crypt depth (μm) | 119.20<sup>a</sup> | 126.76<sup>a</sup> | 89.47<sup>b</sup> | 85.61<sup>b</sup> | 91.96<sup>b</sup> | 89.44<sup>b</sup> | 90.06<sup>b</sup> | 91.51<sup>b</sup> | 3.86 | 0.01 |
| villus height/crypt depth | 4.22<sup>B</sup> | 3.31<sup>B</sup> | 9.47<sup>A</sup> | 9.94<sup>A</sup> | 8.60<sup>A</sup> | 9.29<sup>A</sup> | 9.00<sup>A</sup> | 8.81<sup>A</sup> | 0.53 | < 0.01 |
| 42d |  |  |  |  |  |  |  |  |  |  |
| jejunum |  |  |  |  |  |  |  |  |  |  |
| villus height (μm) | 802.48<sup>b</sup> | 698.15<sup>b</sup> | 1133.07<sup>a</sup> | 1159.25<sup>a</sup> | 1081.09<sup>a</sup> | 1188.26<sup>a</sup> | 1131.01<sup>a</sup> | 1107.43<sup>a</sup> | 44.88 | 0.01 |
| crypt depth (μm) | 123.15<sup>a</sup> | 137.60<sup>a</sup> | 81.18<sup>b</sup> | 80.67<sup>b</sup> | 87.15<sup>b</sup> | 80.34<sup>b</sup> | 84.06<sup>b</sup> | 85.72<sup>b</sup> | 5.46 | 0.01 |
| villus height/crypt depth | 6.85<sup>BCb</sup> | 5.28<sup>C</sup> | 14.69<sup>A</sup> | 14.41<sup>A</sup> | 12.59<sup>ABa</sup> | 14.92<sup>A</sup> | 13.46<sup>ABa</sup> | 13.18<sup>ABa</sup> | 0.87 | < 0.01 |
| ileal |  |  |  |  |  |  |  |  |  |  |
| villus height (μm) | 648.73<sup>BCb</sup> | 571.34<sup>C</sup> | 1249.72<sup>A</sup> | 1273.86<sup>A</sup> | 988.50<sup>ABa</sup> | 1271.14<sup>A</sup> | 1187.20<sup>A</sup> | 993.01<sup>ABa</sup> | 60.20 | < 0.01 |
| crypt depth (μm) | 130.43<sup>A</sup> | 144.66<sup>A</sup> | 77.77<sup>B</sup> | 73.67<sup>B</sup> | 89.01<sup>B</sup> | 76.08<sup>B</sup> | 78.54<sup>B</sup> | 83.33<sup>B</sup> | 5.53 | < 0.01 |
| villus height/crypt depth | 5.01<sup>D</sup> | 4.01<sup>D</sup> | 16.11<sup>A</sup> | 17.33<sup>A</sup> | 11.08<sup>C</sup> | 16.62<sup>A</sup> | 15.06<sup>ABa</sup> | 11.91<sup>BCb</sup> | 1.04 | < 0.01 |

**Fig 1.**
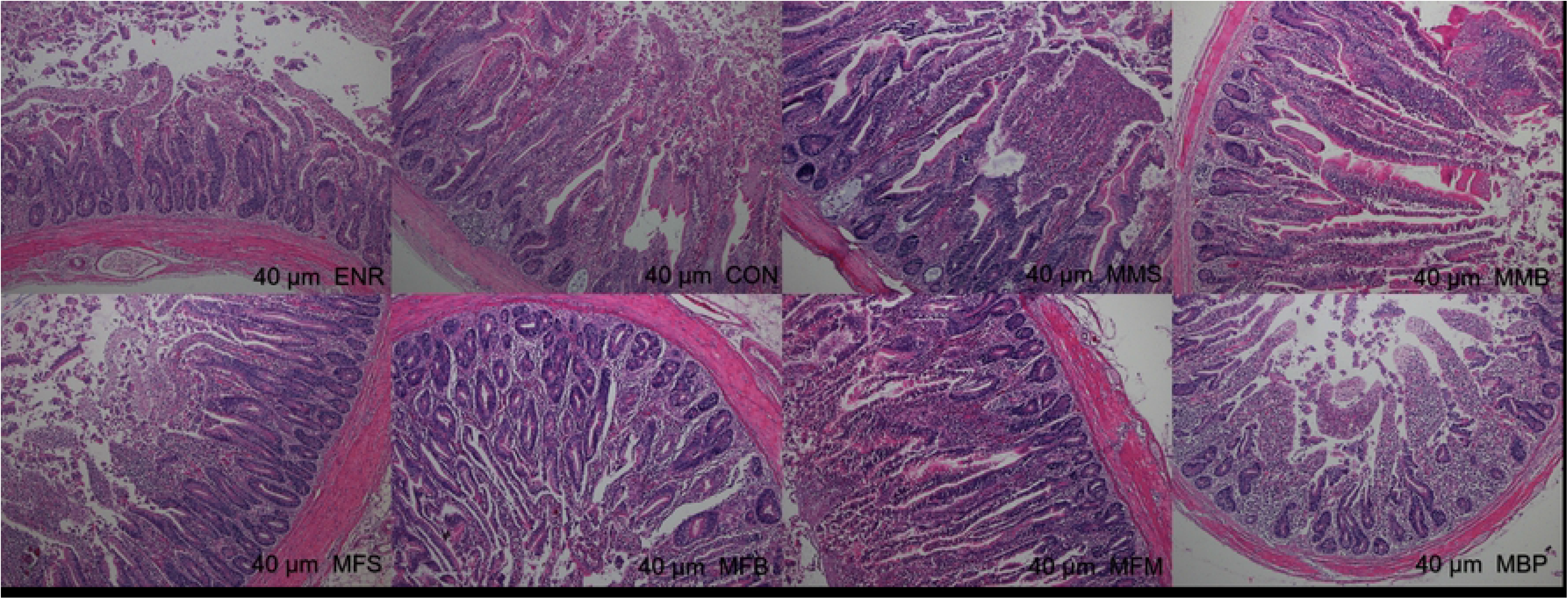
Histological structure of Hematoxylin and Eosin (H&E) staining jejunum at 21 day of age. CON, control diet; ENR; CON + 100 mg/kg ENR, MMS; CON + 2000 mg/kg MOS + 300 mg/kg MAN + 1500 mg/kg SB, MMB; CON + 2000 mg/kg MOS + 300 mg/kg MAN + 500 mg/kg BS, MFS; CON + 2000 mg/kg MOS + 9000 mg/kg FOS + 1500 mg/kg SB, MFB; CON + 2000 mg/kg MOS + 9000 mg/kg FOS + 500 mg/kg BS, MFM; CON + 2000 mg/kg MOS + 9000 mg/kg FOS + 300 mg/kg MAN, MBP; CON + 2000 mg/kg MOS + 500 mg/kg BS + 37 mg/kg PT. SB; Sodium butyrate, FOS; Fructose oligosaccharide, PT; Phytase, BS; Bacillus subtilis, MOS; Mannose oligosaccharide, MAN; Mannanase.

**Fig 2.**
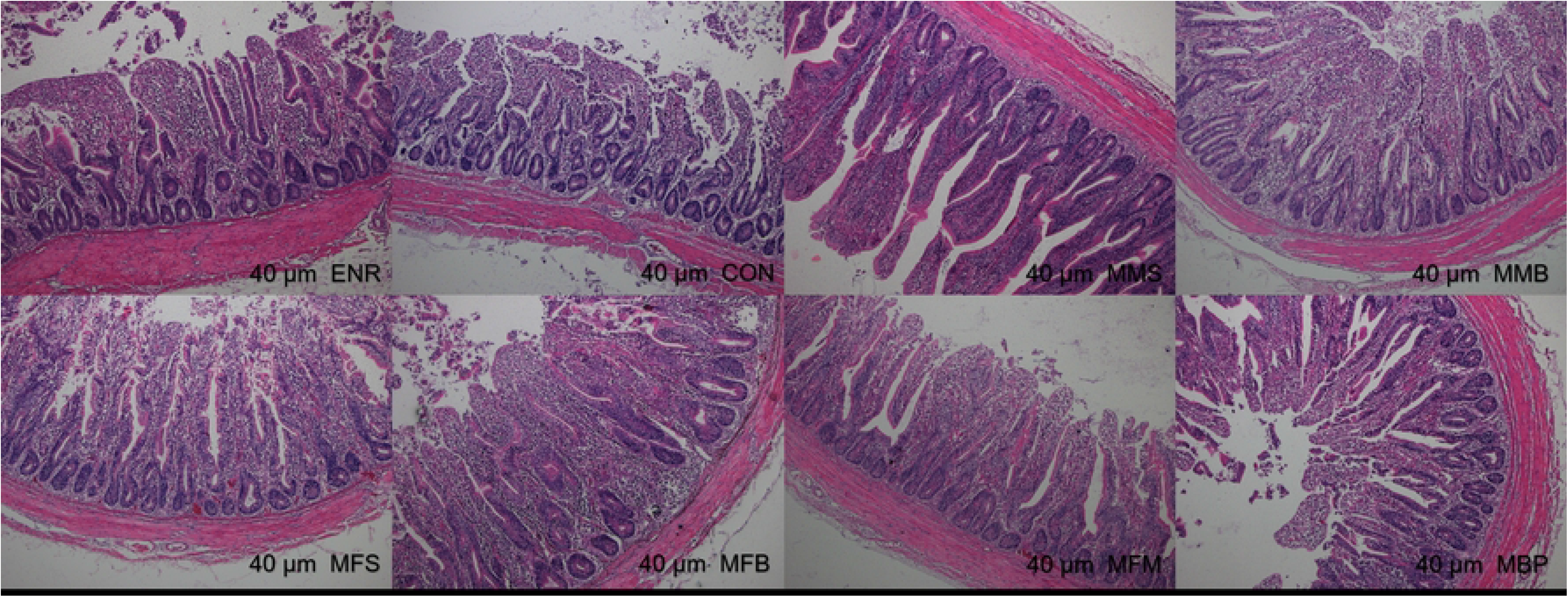
Histological structure of Hematoxylin and Eosin (H&E) staining ileum at 21 day of age. CON, control diet; ENR; CON + 100 mg/kg ENR, MMS; CON + 2000 mg/kg MOS + 300 mg/kg MAN + 1500 mg/kg SB, MMB; CON + 2000 mg/kg MOS + 300 mg/kg MAN + 500 mg/kg BS, MFS; CON + 2000 mg/kg MOS + 9000 mg/kg FOS + 1500 mg/kg SB, MFB; CON + 2000 mg/kg MOS + 9000 mg/kg FOS + 500 mg/kg BS, MFM; CON + 2000 mg/kg MOS + 9000 mg/kg FOS + 300 mg/kg MAN, MBP; CON + 2000 mg/kg MOS + 500 mg/kg BS + 37 mg/kg PT. SB; Sodium butyrate, FOS; Fructose oligosaccharide, PT; Phytase, BS; Bacillus subtilis, MOS; Mannose oligosaccharide, MAN; Mannanase.

**Fig 3.**
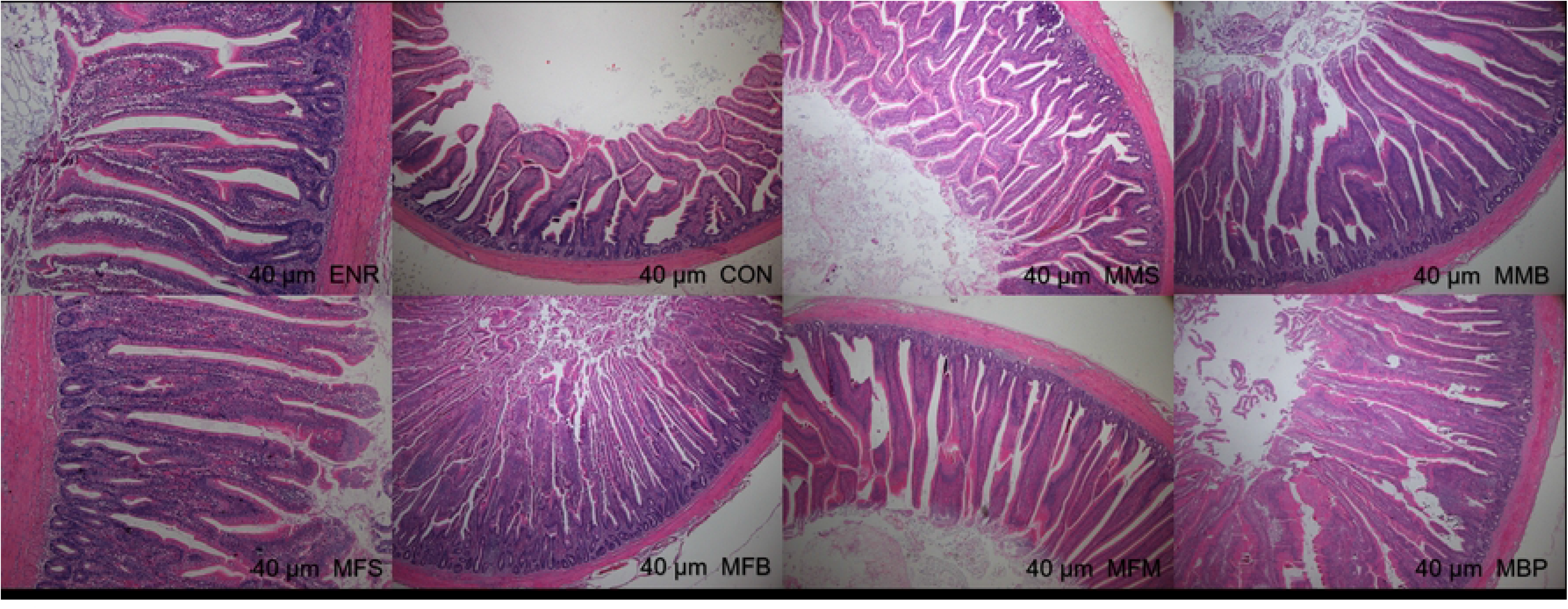
Histological structure of Hematoxylin and Eosin (H&E) staining jejunum at 42 day of age. CON, control diet; ENR; CON + 100 mg/kg ENR, MMS; CON + 2000 mg/kg MOS + 300 mg/kg MAN + 1500 mg/kg SB, MMB; CON + 2000 mg/kg MOS + 300 mg/kg MAN + 500 mg/kg BS, MFS; CON + 2000 mg/kg MOS + 9000 mg/kg FOS + 1500 mg/kg SB, MFB; CON + 2000 mg/kg MOS + 9000 mg/kg FOS + 500 mg/kg BS, MFM; CON + 2000 mg/kg MOS + 9000 mg/kg FOS + 300 mg/kg MAN, MBP; CON + 2000 mg/kg MOS + 500 mg/kg BS + 37 mg/kg PT. SB; Sodium butyrate, FOS; Fructose oligosaccharide, PT; Phytase, BS; Bacillus subtilis, MOS; Mannose oligosaccharide, MAN; Mannanase.

**Fig 4.**
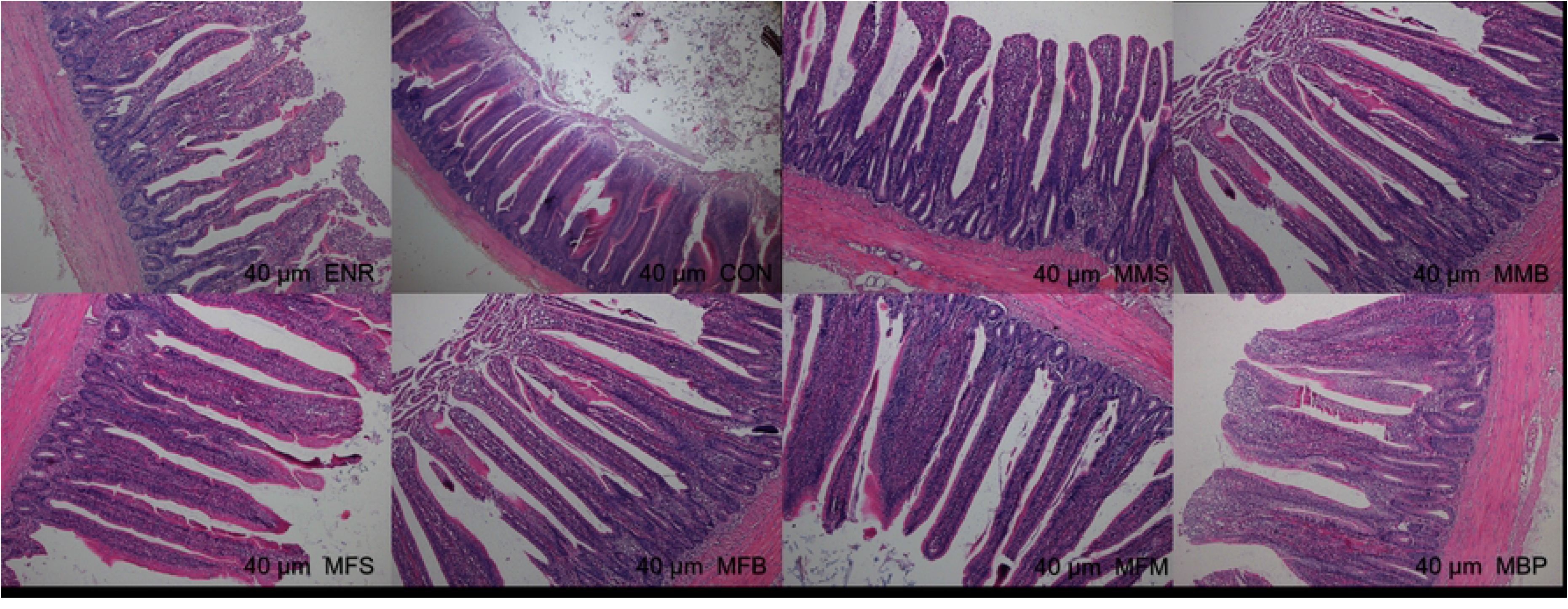
Histological structure of Hematoxylin and Eosin (H&E) staining ileum at 42 day of age. CON, control diet; ENR; CON + 100 mg/kg ENR, MMS; CON + 2000 mg/kg MOS + 300 mg/kg MAN + 1500 mg/kg SB, MMB; CON + 2000 mg/kg MOS + 300 mg/kg MAN + 500 mg/kg BS, MFS; CON + 2000 mg/kg MOS + 9000 mg/kg FOS + 1500 mg/kg SB, MFB; CON + 2000 mg/kg MOS + 9000 mg/kg FOS + 500 mg/kg BS, MFM; CON + 2000 mg/kg MOS + 9000 mg/kg FOS + 300 mg/kg MAN, MBP; CON + 2000 mg/kg MOS + 500 mg/kg BS + 37 mg/kg PT. SB; Sodium butyrate, FOS; Fructose oligosaccharide, PT; Phytase, BS; Bacillus subtilis, MOS; Mannose oligosaccharide, MAN; Mannanase.

### Composition and differences of cecal microflora

#### Phylum level

Appendix Figs 5A and 5B show the relative abundance of the top 10 microorganisms at the phylum level in 21- and 42-d-old birds. The cecal microbiome of each group at 21 d was dominated by *Bacteroidetes*, *Firmicutes*, *Proteobacteria*, *Tenericutes*, *Verrucomicrobia*, *Actinobacteria*, *Cyanobacteria*, *Acidobacteria*, *Fusobacteria*, and *Epsilonbacteraeota*. The cecal microbiome of each group at 42 d of age was dominated by *Bacteroidetes*, *Firmicutes*, *Proteobacteria*, *Tenericutes*, *Lentisphaerae*, *Actinobacteria*, *Cyanobacteria*, *Acidobacteria*, *Fusobacteria*, and *Epsilonbacteraeota*. Among these, *Bacteroidetes* and *Firmicutes* were the most dominant bacterial groups, which together accounted for more than 80% of the total microbial community detected. Appendix Figs 5C and 5D show compared with the ENR and CON groups, the MMS, MMB, and MBP groups showed increase in the abundance of *Firmicutes* at 21 d and in the abundance of *Bacteroides* at 42 d, whereas the MMB, MFB, and MBP groups showed decreased abundance of *Proteobacteria* at 21 and 42 d.

**Fig 5.**
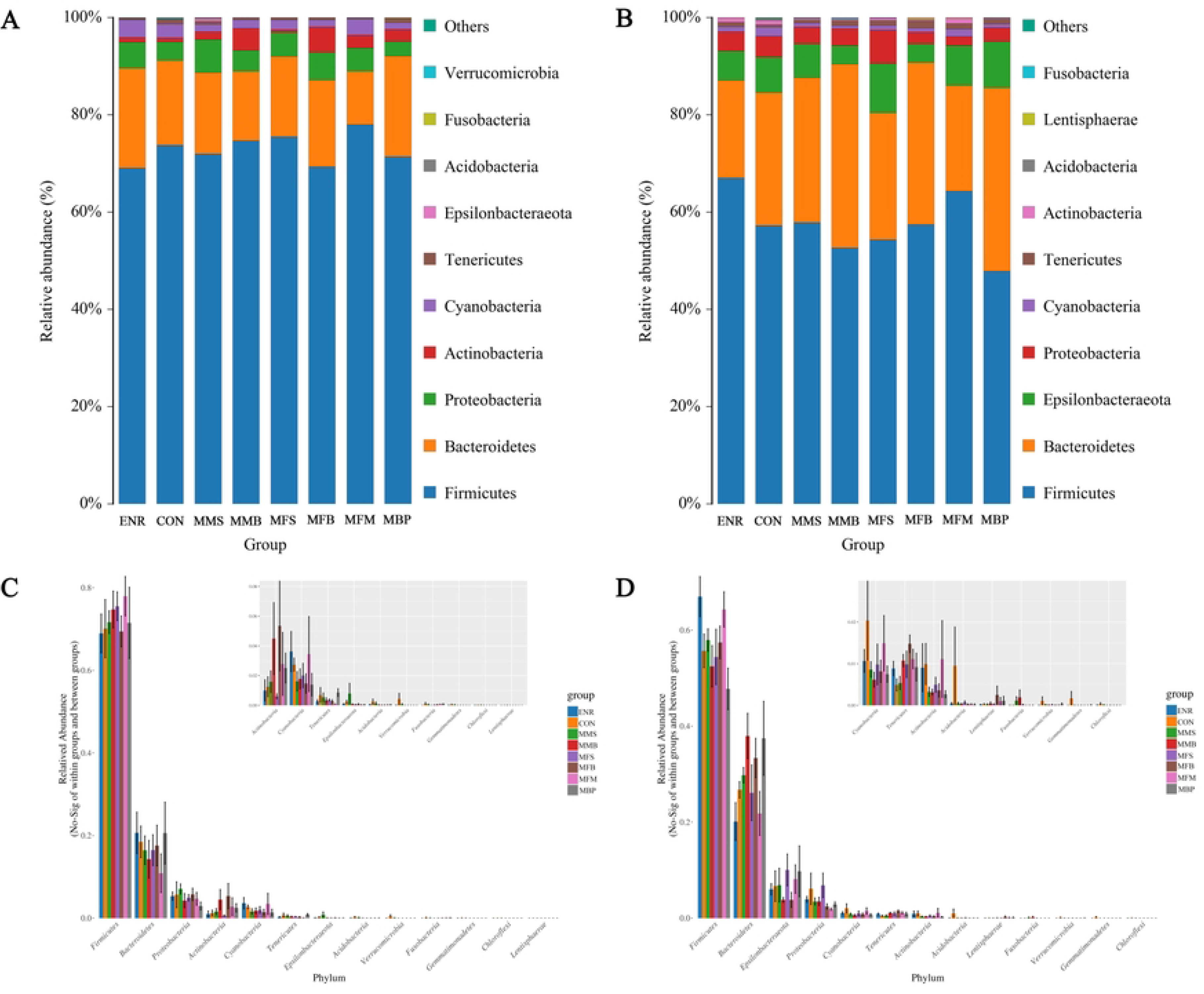
Different taxa and significantly different taxa between different groups by Anova analysis at the phylum level at 21 and 42 days old. (A) and (C) 21d. (B) and (D) 42d. CON, control diet; ENR; CON + 100 mg/kg ENR, MMS; CON + 2000 mg/kg MOS + 300 mg/kg MAN + 1500 mg/kg SB, MMB; CON + 2000 mg/kg MOS + 300 mg/kg MAN + 500 mg/kg BS, MFS; CON + 2000 mg/kg MOS + 9000 mg/kg FOS + 1500 mg/kg SB, MFB; CON + 2000 mg/kg MOS + 9000 mg/kg FOS + 500 mg/kg BS, MFM; CON + 2000 mg/kg MOS + 9000 mg/kg FOS + 300 mg/kg MAN, MBP; CON + 2000 mg/kg MOS + 500 mg/kg BS + 37 mg/kg PT. SB; Sodium butyrate, FOS; Fructose oligosaccharide, PT; Phytase, BS; Bacillus subtilis, MOS; Mannose oligosaccharide, MAN; Mannanase.

#### Genus level

Appendix Figs 6A and 6B show the relative abundance of the top 10 microorganisms at the genus level in 21- and 42-d-old birds. Appendix Figs 6C and 6D show compared with that in the CON and ENR groups, the relative abundance of *Sellimonas* in the other groups was increased and that of *Pseudochrobactrum* was decreased at 21 d. Compared with that in the CON group, the relative abundance of *UCG-013* in *Ruminococcaceae* was increased at 42 d in the other groups.

**Fig 6.**
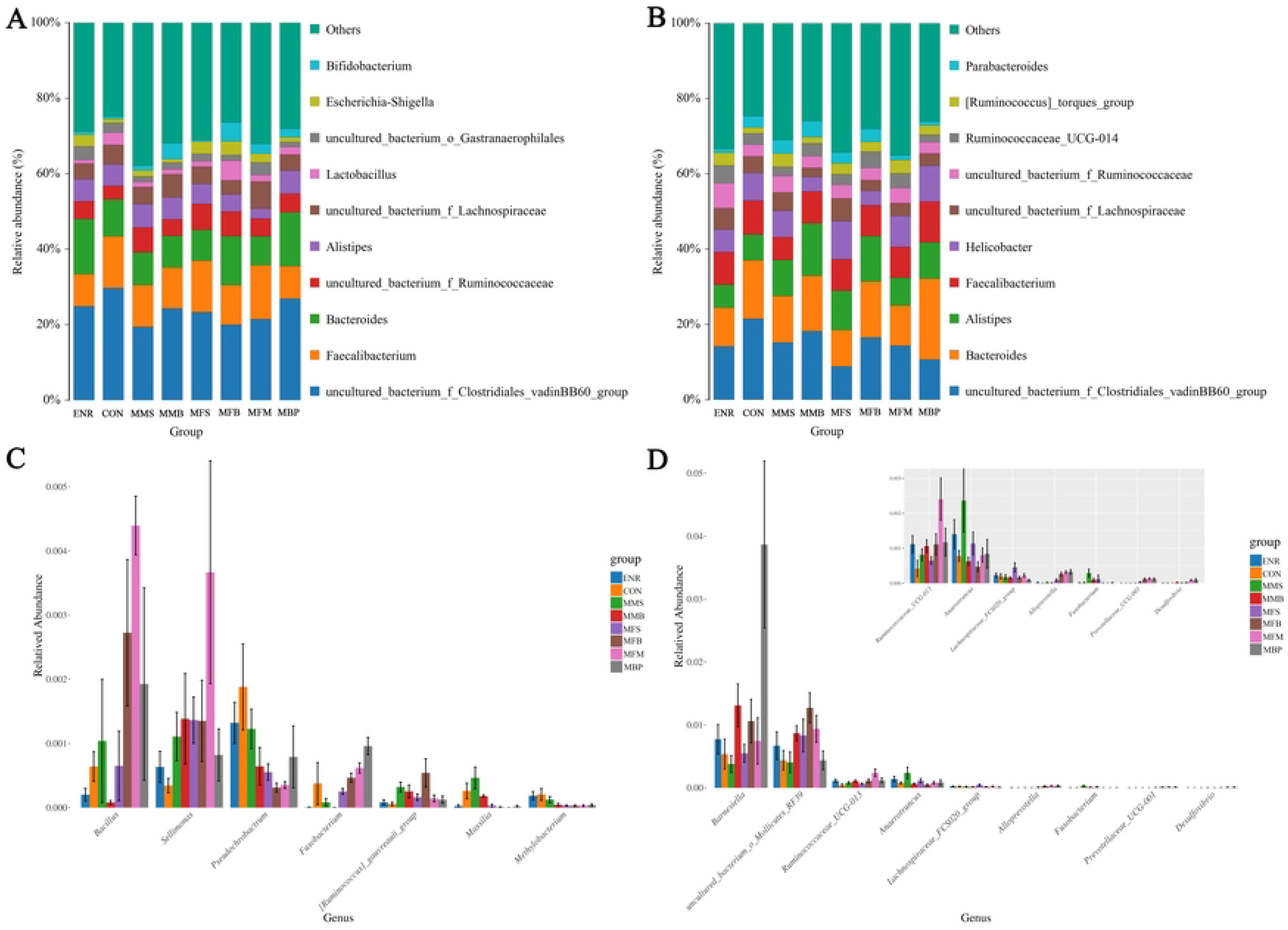
Different taxa and significantly different taxa between different groups by Anova analysis at the genus level at 21 and 42 days old. (A) and (C) 21d. (B) and (D) 42d. CON, control diet; ENR; CON + 100 mg/kg ENR, MMS; CON + 2000 mg/kg MOS + 300 mg/kg MAN + 1500 mg/kg SB, MMB; CON + 2000 mg/kg MOS + 300 mg/kg MAN + 500 mg/kg BS, MFS; CON + 2000 mg/kg MOS + 9000 mg/kg FOS + 1500 mg/kg SB, MFB; CON + 2000 mg/kg MOS + 9000 mg/kg FOS + 500 mg/kg BS, MFM; CON + 2000 mg/kg MOS + 9000 mg/kg FOS + 300 mg/kg MAN, MBP; CON + 2000 mg/kg MOS + 500 mg/kg BS + 37 mg/kg PT. SB; Sodium butyrate, FOS; Fructose oligosaccharide, PT; Phytase, BS; Bacillus subtilis, MOS; Mannose oligosaccharide, MAN; Mannanase.

### Diversity of cecal microbiota

#### Alpha diversity

Alpha diversity among the ENR, CON, MMS, MMB, MFS, MFB, MFM, and MBP groups is presented in Table 5. The alpha diversity indexes were calculated based on the OTUs using the Shannon, Simpson, and Chao1 methods. No significant differences in the alpha diversity indexes (including OTU, Shannon, Simpson, and Chao1) of the cecal microbiota in broilers were observed at 21 and 42 d.

**Table 5.** Effects of nonantibiotic alternative growth promoter combinations on alpha diversity of microbiota in cecum of broilers at 21 and 42 days.

| Items<sup>1</sup> | ENR | CON | MMS | MMB | MFS | MFB | MFM | MBP | SEM | p |
| --- | --- | --- | --- | --- | --- | --- | --- | --- | --- | --- |
| 21d |  |  |  |  |  |  |  |  |  |  |
| Feature | 364.17 | 353.17 | 371.33 | 345.17 | 379.67 | 351.67 | 351.67 | 355.67 | 4.32 | 0.52 |
| Chao1 | 402.3 | 385.17 | 398.25 | 400.06 | 418.72 | 411.59 | 394.53 | 388.43 | 4.66 | 0.68 |
| Simpson | 0.951 | 0.948 | 0.96 | 0.955 | 0.952 | 0.931 | 0.935 | 0.949 | 0.004 | 0.49 |
| Shannon | 5.64 | 5.54 | 5.94 | 5.66 | 5.75 | 5.28 | 5.52 | 5.8 | 0.06 | 0.21 |
| 42d |  |  |  |  |  |  |  |  |  |  |
| Feature | 431.67 | 389.83 | 444.83 | 453.5 | 448.67 | 451 | 437 | 418.83 | 6.52 | 0.22 |
| Chao1 | 465.24 | 436.69 | 478.43 | 488.43 | 477.73 | 480.8 | 475.04 | 454.34 | 5.8 | 0.4 |
| Simpson | 0.964 | 0.95 | 0.964 | 0.966 | 0.956 | 0.967 | 0.957 | 0.942 | 0.003 | 0.28 |
| Shannon | 6.19 | 5.89 | 6.23 | 6.15 | 6.07 | 6.26 | 6.06 | 5.45 | 0.08 | 0.14 |

<sup>1</sup>CON, control diet; ENR; CON + 100 mg/kg ENR, MMS; CON + 2000 mg/kg MOS + 300 mg/kg MAN + 1500 mg/kg SB, MMB; CON + 2000 mg/kg MOS + 300 mg/kg MAN + 500 mg/kg BS, MFS; CON + 2000 mg/kg MOS + 9000 mg/kg FOS + 1500 mg/kg SB, MFB; CON + 2000 mg/kg MOS + 9000 mg/kg FOS + 500 mg/kg BS, MFM; CON + 2000 mg/kg MOS + 9000 mg/kg FOS + 300 mg/kg MAN, MBP; CON + 2000 mg/kg MOS + 500 mg/kg BS + 37 mg/kg PT. SB; Sodium butyrate, FOS; Fructose oligosaccharide, PT; Phytase, BS; Bacillus subtilis, MOS; Mannose oligosaccharide, MAN; Mannanase.

#### Beta diversity

Beta diversity analysis was performed to compare the degree of similarity among different samples with respect to species. Beta diversity was assessed by PCoA using the weighted UniFrac distance method. Figs 7A and 7B show the PCoA results of the variation among the eight groups at 21 and 42 d; no significant difference in species diversity was observed. Linear discriminant analysis effect size (LEfSe) analysis was used to determine biomarkers with significant differences in expression among the different treatments. Figs 7C and 7D show the species with significant differences among the eight groups at 21 and 42 d with LDA scores > 4. *Ruminiclostridium* at 21d as well as *Tannerellaceae* and *Parabacteroides* at 42d showed LDA scores > 4 in the MMS group.

**Fig 7.**
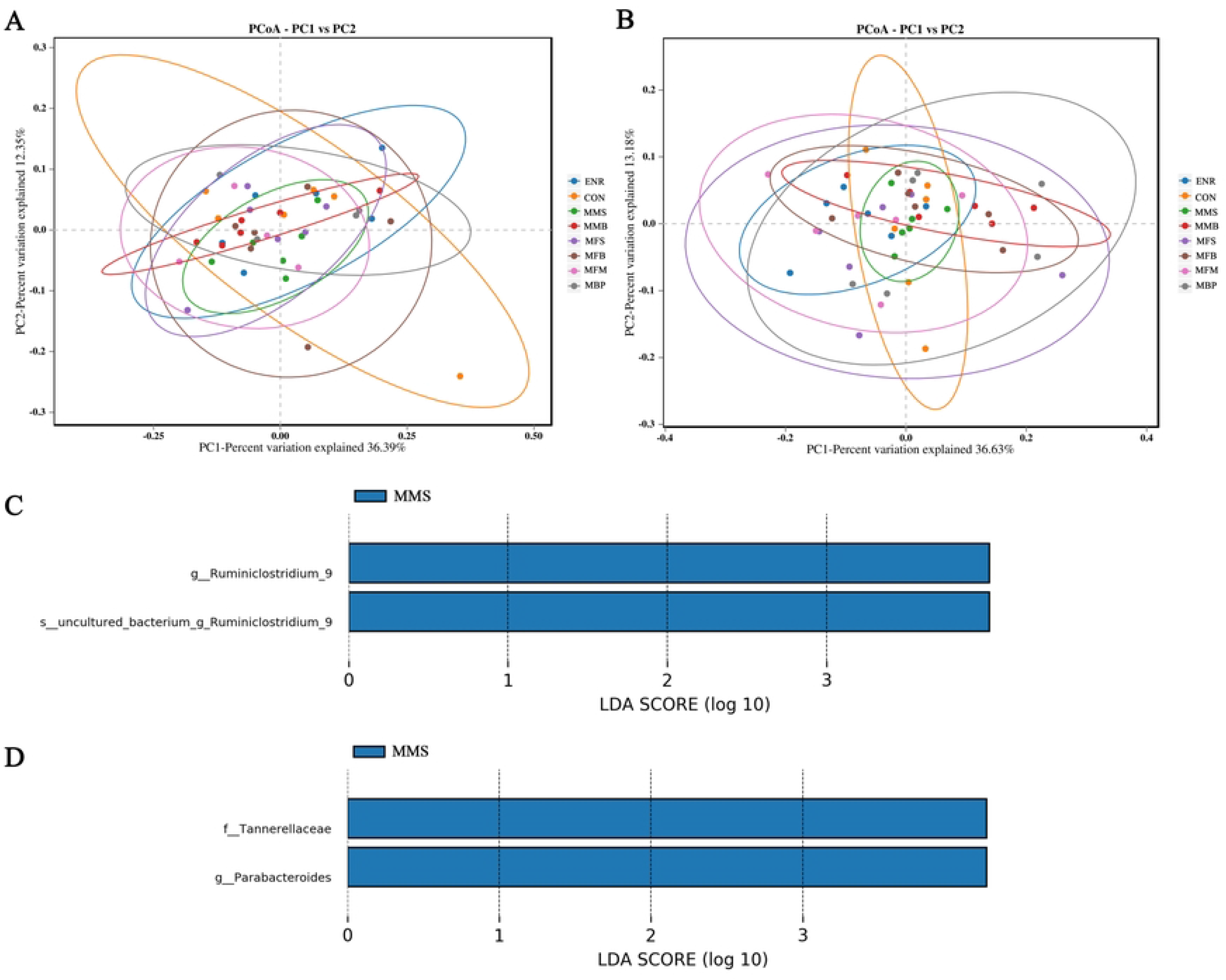
Beta diversity and Linear discriminant analysis effect size analysis of the microbiome residing in the cecal chyme of broilers at 21 and 42 days old. (A) PCoA plot at 21d. (B) PCoA plot at 42d. (C) LDA distribution histogram at 21d (LDA scores > 4). (C) LDA distribution histogram at 42d (LDA scores > 4). CON, control diet; ENR; CON + 100 mg/kg ENR, MMS; CON + 2000 mg/kg MOS + 300 mg/kg MAN + 1500 mg/kg SB, MMB; CON + 2000 mg/kg MOS + 300 mg/kg MAN + 500 mg/kg BS, MFS; CON + 2000 mg/kg MOS + 9000 mg/kg FOS + 1500 mg/kg SB, MFB; CON + 2000 mg/kg MOS + 9000 mg/kg FOS + 500 mg/kg BS, MFM; CON + 2000 mg/kg MOS + 9000 mg/kg FOS + 300 mg/kg MAN, MBP; CON + 2000 mg/kg MOS + 500 mg/kg BS + 37 mg/kg PT. SB; Sodium butyrate, FOS; Fructose oligosaccharide, PT; Phytase, BS; Bacillus subtilis, MOS; Mannose oligosaccharide, MAN; Mannanase.

## Discussion

Due to the absence of AGPs, alternate methods need to be developed to ensure feed efficiency and broiler health, and now NAGPCs that combine the same kinds of NAGPs in broilers may limit the beneficial effect of synergy. The aim of the current experiment was to evaluate the effects of six NAGPCs with different types of NAGPs on nutrient utilization, digestive enzyme activity, intestinal morphology, and cecal microflora of broilers so as to find a better alternative method to ensure feed efficiency and the broiler health. We found that NAGPC supplementation significantly increased the utilization of DM, OM, CP, and CF at 21 and 42 d as well as the utilization of CA and P at 42 d. The synergistic and complementary effects of additives observed in this study may be attributed to the different beneficial mechanisms of action of each additive [16]. Previous studies have reported that dietary supplementation of MAN [17, 18] and PT [19] results in improved nutritional digestibility in broilers, possibly because of the beneficial effect of MAN in reducing the action of mannan, improving nutrient absorption by the release of encapsulated nutrients through breakdown of the cell wall matrix, and decreasing digesta viscosity [20]. The benefits of PT supplementation have been attributed to effects of further phytate degradation [21] and restoration of enzyme functions [22]. Furthermore, prebiotics may disrupt intestinal pathogen colonization and improve the intestinal environment. This is the main reason for improved nutrient utilization [23]. SB has a similar effect, as it helps broilers to reduce the number of pathogens in the digestive tract and regulate intestinal microflora [24, 25]. Studies have shown that adding a combination of prebiotics and butyric acid to the diet of broilers can better control *Salmonella typhimurium* abundance and promote growth compared with addition of prebiotics or butyric acid alone. This could be attributed to the synergistic effect of prebiotics and butyric acid [16]. BS can increase the length of intestinal villi and crypt depth, thus increasing the digestibility of nutrients [26]. Besides, the probiotics use the prebiotics as a food source, which enables them to survive for a longer period of time inside the human digestive system than would otherwise be possible [27]. Finally, the increase in the nutritional utilization rate of birds treated with NAGPCs may be related to the increase in duodenal digestive enzyme activity.

In this study, broilers fed diets supplemented with NAGPCs showed higher activity of trypsin, lipase, and amylase. Some species of pathogenic bacteria, such as *Escherichia coli* [28] and *Clostridium* [29], have been shown to inhibit digestive enzyme secretion by damaging the villi and microvilli in the intestinal mucosa. Organic acids can lower the pH of the chyme, which minimizes the pathogen load and enhances the digestibility of protein by improving pepsin activity [30]. Moreover, they can enhance the production of pancreatic juice containing various zymogens (trypsinogen, chymotrypsinogens A and B, and procarboxypeptidases A and B), leading to improved digestive enzyme activity [31]. Prebiotics are non-digestible carbohydrates that selectively stimulate *Bifidobacterium* and *Lactobacillus* growth [32]; furthermore, the effect of MAN in reducing intestinal viscosity has been suggested to prepare a suitable environment for *Lactobacillus* growth [33]. *Bifidobacterium* and *Lactobacillus* colonizing the intestine have been reported to deliver enzymes [34], explaining why prebiotics and MAN increase the activity of digestive enzymes. Regarding BS, their presence along intestinal sections might stimulate the production of endogenous enzymes by broilers [35]. Furthermore, BS itself can produce protease and amylase [36]. Research has shown that phytate can chelate with the co-factors required for optimum enzyme activity to reduce the activity of digestive enzymes [37] whereas PT can promote the degradation of phytate, thus increasing the activity of digestive enzymes [21]. Finally, the observed synergistic and complementary effect of the additives used in this study can be attributed to the different beneficial mechanisms of action of each NAGP.

Villi and crypts are the two key components of the small intestine, and their shape offers an indication of absorptive ability [38]. Increasing villus height provides an increased surface area for the absorption of available nutrients [39]. The villus crypt is considered the villus factory, and deeper crypts indicate rapid tissue turnover to permit villus renewal as needed in response to normal sloughing or inflammation from pathogens or their toxins and high tissue demands [40, 41]. Intestinal epithelial cells originating in the crypt migrate along the villus surface upward to the villus tip and are extruded into the intestinal lumen within 48 to 96 h [42, 43]. Shortening of the villi and deepening crypts may lead to poor nutrient absorption, increased secretion in the gastrointestinal tract, and lower performance [44]. In contrast, increased villus height and villus height:crypt depth ratio are directly correlated with increased epithelial cell turnover [45], and longer villi are associated with activated cell mitosis [46]. The present results indicated that supplementing NAGPC in broiler diets could improve the villus height and villus height/crypt depth as well as decrease crypt depth in the jejunum and ileum. Previous research indicates that the addition of synbiotics or probiotics can increase the turnover rate of epithelial cells, improve intestinal tissue structure, and increase the ratio of villus height to crypt depth in the ileum [47]. Further, addition of PT to the diet may decline the growth of intestinal pathogenic bacteria by possibly decreasing the quantity of available substrates for their metabolization [48]; this mechanism also reduces the pathogen-induced damage to the intestinal mucosa [49]. MAN can reduce digesta viscosity by degrading β-mannan, one of the major soluble non-starch polysaccharides in the diet, and thus ameliorate structural damage to the absorptive architecture [50]. Finally, SB can be used by intestinal cells to stimulate intestinal development [51] and significantly improve the morphology of the jejunum and ileum [52].

The intestinal microbiota has a significant impact on the control of host homeostasis, organ development, metabolic processes, and immunological response [53, 54]. In this study, the improvement of cecal microflora by NAGPC supplementation may be attributed to the synergistic effect of different NAGPs. Prebiotics are reported to have a beneficial effect on the host by selectively stimulating the growth and/or activity of one or a limited number of bacteria in the intestine or gut [23]. BS can compete with pathogens, balance intestinal microbiota [55]. Organic acids can reduce cecal pH and inhibit the growth of pathogenic bacteria [56] as well as adjust the acidic environment to facilitate *Lactobacillus* survival [57]. The correlation between increased digesta viscosity and increased intestinal concentrations of pathogenic bacteria has previously been reported in broilers [58]; MAN can reduce the viscosity of digestive juice, thus promoting the growth of beneficial microflora [59]. Differences in dietary phosphorus content can also affect the cecal microbial diversity of broilers [48]; phytase can promote the degradation of phytate and release P and other nutrients [60], which may improve the cecal microflora.

In this study, we analyzed the changes in cecal microbial composition at the phylum and genus level. At the phylum level, *Firmicutes* and *Bacteroides* were the dominant bacterial groups, accounting for more than 80% of the total microbial communities detected in this study. *Firmicutes* and *Bacteroides* were shown to be the major phyla of the cecal population in 21 and 42 d broilers in our experiment. This is consistent with prior research, as these bacteria are known to play a role in energy generation and metabolism [61]. The dominating groups of the chosen chickens may alter according to changes in age, breed, and area. According to the current research, *Firmicutes* and *Bacteroides* are the major phyla of the cecal community in 42 d broilers supplemented with dietary BS [62], prebiotics [63], organic acids [64], PT [65], and MAN [33]. Further, *Firmicutes* and *Bacteroides* are involved in carbon metabolism [66] and fat deposition [67] whereas *Proteobacteria* comprises zoonotic disease-causing bacteria such as *Escherichia coli*, *Salmonella*, *Campylobacter*, and other well-known pathogens [68]. The results of our study showed that compared with those in the ENR and CON groups, the abundance of *Firmicutes* at 21 and the abundance of *Bacteroides* at 42 d was increased in the MMS, MMB, and MBP and the abundance of *Proteobacteria* was decreased at 21 and 42 d in the MMB, MFB, and MBP groups.

At the genus level, *Clostridiales*, *Bacteroides*, and *Feacalibacterium* were the dominant genera of the cecal community in 21d broilers in our trial, *Clostridiales*, *Bacteroides*, and *Alistipes* were the dominant genera of the cecal community in 42 d broilers in our trial. *Feacalibacterium* is a butyric acid-generating bacterium found in chicken cecum [69]. *Feacalibacterium* may supply energy to the body and alleviate inflammation; its presence thus signals the host intestinal health [70, 71]. *Alistipes* exert anti-inflammatory properties and may protect against some illnesses [72]. *Clostridiales* are involved in the metabolism of intestinal proteins [73]. *Pseudochrobactrum,* is a potential hazard as a pathogen [74]. The results of our study showed that compared with those in the ENR and CON, NAGPCs addition to the diet could decrease the abundance of *Pseudochrobactrum* at 21d. *Sellimonas* is a potential biomarker of intestinal homeostasis [75], the results show that compared with that in ENR and CON, diets supplemented with NAGPCs could increase the abundance of *Sellimonas* at 21d.

The diversity of intestinal microflora is critical for maintaining the gastrointestinal equilibrium and is helpful to the host health [54]. In our study, dietary treatments failed to modify the overall diversity of cecal microbiota at 21 and 42 d. These results were consistent with previous reports. Current research indicates that supplementation with BS [76], FOS, MOS [23], SB [77], MAN [78], and PT [79] has no effect on the diversity and community structure of cecal microbiota in broilers. In fact, aging has a greater impact on microbiota than that of therapy [80]. *Ruminiclostridium* at 21d and *Tannerellaceae* and *Parabacteroides* at 42d with LDA scores > 4 were observed in the MMS group. *Ruminiclostridium*, as a beneficial bacterium, can secrete cellulase and related plant cell wall-degrading enzymes, thereby degrading cellulose in the feed and releasing nutrients [81]. *Tannerellaceae* can secrete butyric acid and propionic acid to regulate cecal pH [82]. *Parabacteroides* have the physiological characteristics of carbohydrate metabolism and secrete short chain fatty acids [83]. This may be caused by the synergistic effect of MAN, MOS, and SB.

As legislation and public demand for “antibiotic-free” poultry increase the pressure to abandon the use of AGPs, other techniques to stimulate broiler chicken development are being explored. The present research compared six NAGPCs to evaluate their effectiveness as growth promoters and to analyze indicators of broiler health. The results showed that compared with the CON and ENR groups, MMS, MMB, MFS, MFB, MFM, and MBP groups showed significantly improved nutrient utilization. Duodenal digestive enzyme activity and ileal and jejunal histology are promising biomarkers for explaining these nutrient utilization effects. However, the specific mechanisms of action need identified for successfully replacing antibiotic growth promoters. Overall, ideal combinations of different alternatives are likely to be the key to increasing broiler performance and preserving productivity.

## Acknowledgments

This work was financially supported by Shandong Province Nonantibiotic Alternative Growth Promoter Combinations and Creation of Feed Without Antibiotic products Project (2019JZZY020609-02), Technological system of Modern Agricultural Industry in Shandong Province Sheep Industry Innovation team (SDAIT-10-04), Qingdao Agricultural University Doctoral Start-Up Fund (663/1120009).

